# CHOLINERGIC MODULATION OF CELLULAR RESONANCE IN NON-HUMAN PRIMATE HIPPOCAMPUS

**DOI:** 10.1101/2025.01.10.632495

**Authors:** Abigail Gambrill, Jon W. Rueckemann, Andres Barria

## Abstract

Acetylcholine modulates the network physiology of the hippocampus, a crucial brain structure that supports cognition and memory formation in mammals ^1–3^. In this and adjacent regions, synchronized neuronal activity within theta-band oscillations (4-10Hz) is correlated with attentive processing that leads to successful memory encoding ^4–10^. Acetylcholine facilitates the hippocampus entering a theta oscillatory regime and modulates the temporal organization of activity within theta oscillations ^11,12^.

Unlike rodents that exhibit constant theta oscillations during movement and exploration, primates only manifest theta oscillations in transient bouts during periods of acute attention—despite conserved hippocampal anatomy ^13–16^. The phasic nature of primate theta oscillations and their susceptibility to muscarinic antagonists ^17^, suggest that acetylcholine afferents acutely modulate local circuitry, resulting in a temporary shift in hippocampal rhythmic dynamics. However, we lack a mechanistic understanding that links cellular physiology to emergent theta-rhythmic network dynamics.

We explored the hypothesis that acetylcholine induces a distinct modulation of cellular properties to facilitate synchronization within the theta band in non-human primate neurons.

Here we show that non-human primate neurons from the CA1 region of monkey hippocampus are not homogeneous in their voltage response to inputs of varying frequencies, a phenomenon known as cellular resonance ^18,19^. We classified these neurons as ‘resonant’ or ‘non-resonant’. Under the influence of carbachol, these two classes of neurons become indistinguishable in their resonance, suggesting that acetylcholine transiently creates a homogeneous susceptibility to inputs within the theta range. This change is mediated by metabotropic acetylcholine receptors that enhance sag potentials, indicating that acetylcholine acts on principal neurons to modulate Hyperpolarization-activated Cyclic Nucleotide-gated channels.

Our results reveal a mechanism through which acetylcholine can rapidly modulate intrinsic properties of primate hippocampal neurons to facilitate synchronization within theta-rhythmic circuits, providing insight into the unique features of primate hippocampal physiology.

## Two Classes of Cellular Resonance in the Primate Hippocampus

To characterize the cellular resonance of monkey hippocampal pyramidal neurons, we performed somatic whole-cell recordings at 32℃ on visually identified CA1 neurons in acute slices from the intermediate hippocampus. This sparsely available tissue was obtained through the Tissue Distribution Program of the Washington National Primate Research Center from animals that were transcardially perfused with a neuroprotective N-Methyl-D-glucamine artificial cerebrospinal fluid (NMDG-aCSF) solution during euthanization ^20^. Slices were prepared within 15 minutes of euthanasia and placed in recovery solution (NMDG-aCSF). Neurons were filled with neurobiotin through the recording pipette to verify the identity of monkey CA1 pyramidal cells (Figure 1A).

**Figure 1.**
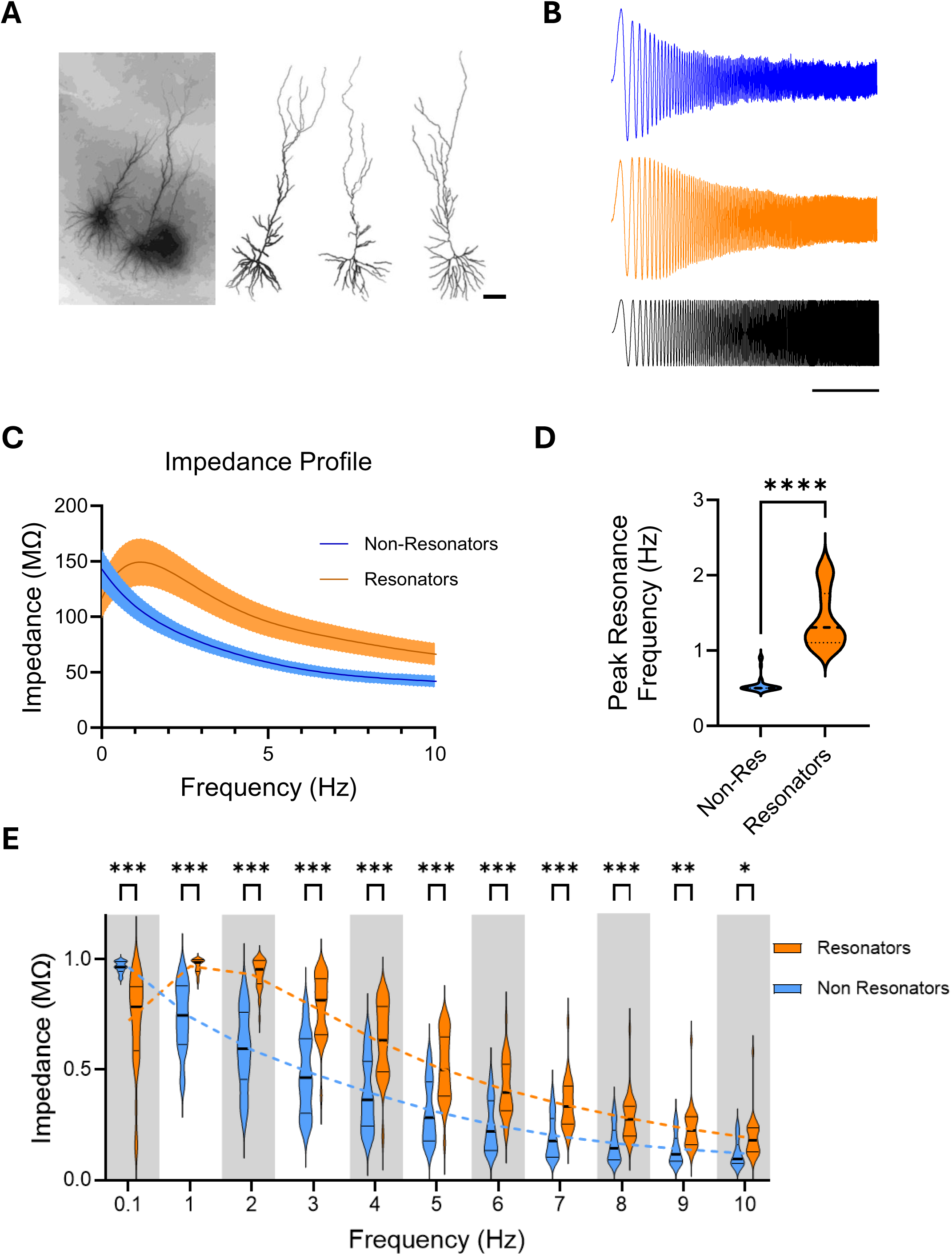
Cellular Resonance of Non-human Primate Hippocampal Neurons. **A.** CA1 sample cells from macaque hippocampal slices filled with neurobiotin during whole-cell recordings and reconstructed post fixation. Scale bar 100 μm. **B.** Sample traces of voltage responses of two types of CA1 hippocampal neurons, non-resonator (top trace) and resonator (middle trace), in response to current injection of constant amplitude and increasing frequency (bottom trace). Scale bar = 5 s. **C.** Average Impedance Profile ± s.e. of non-resonators (n=31) and resonators (n=41) neurons calculated as the ratio of the fast furrier transforms of the voltage response and the injected current. **D.** Preferred Resonant Frequency median (dotted line) and the frequency distribution (width) of the data for non-resonators (blue, n=31) and resonators neurons (orange, n=41). Preferred Resonant Frequency is estimated as the frequency where the impedance is maximal. Statistical significance determined using unpaired Mann-Whitney test (p<0.0001). **E.** Rescaled values of impedance ([0 max]) from non-resonant and resonant cells binned around integer frequencies and compared using a 2-way ANOVA (Frequency x Non-Resonators/Resonators (F(10, 490) = 36.54, P < 0.001)), followed by Sidak’s multiple comparisons test.

Cellular resonance defines the membrane voltage response to incoming stimuli at specific frequencies, capturing the relationship between input rhythmicity and neuronal excitability ^18^. It was determined by measuring voltage responses to subthreshold sinusoidal currents of constant amplitude across varying frequencies (Figure 1B). For these measurements, cells were current-clamped to allow fluctuations in membrane voltage from a baseline of −70 mV.

Cellular resonance is further characterized by the impedance profile ^21^, a key indicator that identifies the input frequency at which the impedance—and consequently the voltage response— reaches its maximum, a point termed the Preferred Resonance Frequency. The impedance profile is calculated from the ratio of the Fast Fourier Transform (FFT) spectrum of the voltage response to that of the injected current.

Analysis across all recorded cells revealed two distinct groups of neurons with different Preferred Resonance Frequency values (Supplemental Figure 1A and B). We classified cells with impedance profiles resembling pink noise filtering, i.e., a response like a 1/f function with a Preferred Resonance Frequency less than 1 Hz, as ‘non-resonators.’ In contrast, neurons that exhibited lower impedance at lower frequencies and increased impedance at higher frequencies—which amplified voltage responses near the frequency of theta-band oscillations—were designated as ‘resonators’ (Figure 1C). Cellular resonance profiles were readily separable by the Preferred Resonance Frequency (Figure 1D).

Somatic recordings at 32°C provided better stability for the slices but yielded lower Preferred Resonance Frequency values compared to extracellularly recorded theta oscillations. Further recordings in a subset of neurons at in vivo brain temperature (38°C) confirmed that Preferred Resonance Frequency is temperature-dependent, with higher temperatures yielding frequencies closer to those of observed extracellular theta oscillations (Supplemental Figure 1C).

In addition to differences in Preferred Resonant Frequency, resonators differed from non-resonators by having attenuated voltage responses at lower frequencies and amplified voltage responses across a range of higher frequencies. An analysis of normalized binned impedance profile values was conducted using a two-way ANOVA, which revealed a significant interaction between Frequency and Non-Resonators/Resonators (F(10, 490) = 36.54, P < 0.001). This interaction indicates that the difference between Resonator and Non-Resonator cells depends on frequency. Post-hoc comparisons using Šídák’s multiple comparisons test confirmed significant differences between Resonator and Non-Resonator cells at each frequency level (P < 0.01) (Figure 1E). The pink noise filtering profile of non-resonators suggests that their membrane voltage response is dominated by ‘passive,’ capacitive properties. In contrast, the shift toward higher frequencies in resonators is consistent with increased recruitment of voltage-dependent conductances, or ‘active properties,’ that elicit voltage rebound and are constrained by channel kinetics ^18^. Convergently, the average Preferred Resonance Frequency increased with increased hyperpolarization of the baseline voltage (Supplemental Figure 1D), reflective of recruitment of voltage-gated channels in resonance ^22^.

The difference in resonance profiles across neuron classes was not due to variations in intrinsic passive biophysical properties, such as membrane resistance or resting membrane potential, as these parameters did not differ between the two classes (Supplemental Figure 2A and B). Within these groups, although the membrane resistance positively correlates with the maximum impedance observed, as anticipated, the Preferred Resonance Frequency is not predicted by the membrane resistance (Supplemental Figure 2C and D). This supports the idea that differences in Preferred Resonance Frequency are not due to different recording or cellular conditions. Similarly, maximum impedance does not correlate with the preferred resonance frequency (Supplemental Figure 2E).

The ability of neurons to produce an enhanced voltage response to some preferred frequency of input is believed to be instrumental in facilitating the synchronization of neurons within a microcircuit, allowing oscillation of the ensemble within the theta band frequency ^23^. The heterogeneity observed in the resonance capacity of hippocampal neurons suggests that neuromodulatory mechanisms may regulate their resonance perhaps to facilitate their incorporation into oscillatory micro circuits engaged in a cognitive event.

## Cholinergic Modulation of Cellular Resonance

Acetylcholine supports a slower-frequency (Type II) theta oscillation in rodents that transiently occurs during events that elicit acute attention, such as receiving reward or encountering novel stimuli ^12^. In the hippocampus of primates, similar slower-frequency theta oscillations are observed in short-lived bouts^13–16^. The phasic nature of theta in primates and its similarity to rodent Type II theta suggests that acetylcholine afferents may have an analogous role in supporting theta activity in the primate hippocampus.

To develop a mechanistic understanding of the role of acetylcholine in the primate hippocampus, we examined the effect of carbachol, a cholinergic receptor agonist, on the resonance properties of monkey hippocampal CA1 neurons. The voltage response to a sinusoidal injected current of varying frequency was recorded in the same cell before and 10 minutes after the addition of carbachol to the perfusion bath.

Carbachol rapidly modified the voltage responses of non-resonator neurons (Figure 2A) but did not alter the voltage response of resonator neurons (Figure 2B). Consequently, the impedance profile of the non-resonator neurons was altered (Figure 2C), attenuating the voltage response at lower frequencies and amplifying it at higher frequencies, effectively tuning the band of frequency filtering (Supplemental Figure 3A). This led to a significant increase in the preferred resonance frequency (Figure 2D) and Q, a dimensionless parameter that measures the amplification of preferred frequencies (Figure 2E). Q is calculated as the ratio of the Preferred Resonant Frequency to the range of frequencies where the impedance’s amplitude is at least half of its maximum value. In contrast, carbachol did not affect the impedance profile, the preferred resonance frequency, or the Q parameter of resonator neurons (Figures 2F-H and Supplemental Figure 3B).

**Figure 2.**
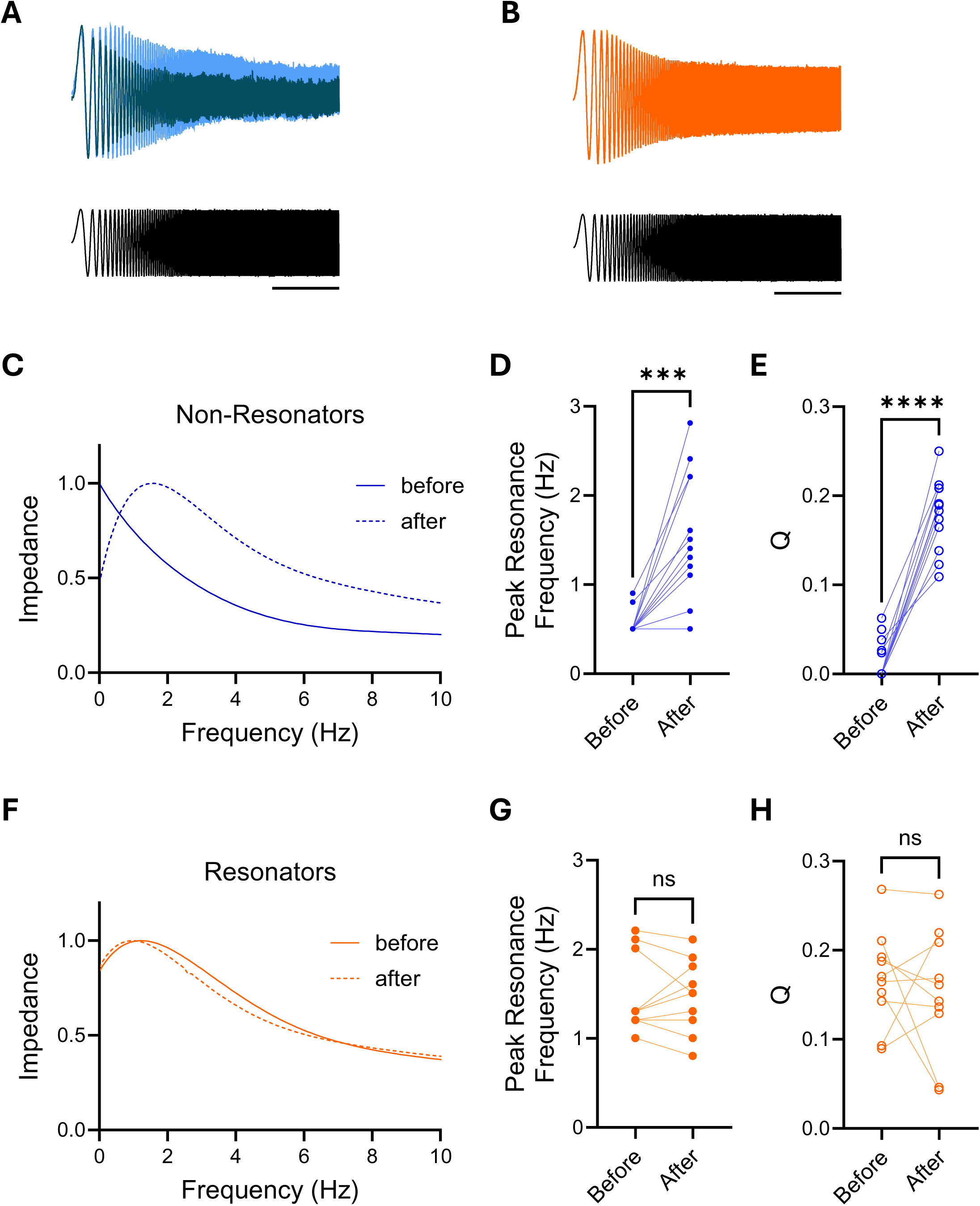
Effect of Carbachol on Cellular Resonance. **A.** Superimposed voltage response traces from the same non-resonator neuron in response to current injection of constant amplitude and increasing frequency before (dark blue) and 10 minutes after carbachol is added to the bath (light blue). Scale bar = 5 s. **B.** Superimposed voltage response traces of the same resonator neuron in response to current injection of constant amplitude and increasing frequency before (dark orange) and 10 minutes after carbachol is added to the bath (light orange). Scale bar = 5 s. **C.** Sample impedance profile of a non-resonator cell before (continuous line) and after (dotted line) carbachol has been added to the bath. **D.** Peak Resonance Frequency of non-resonant cells before and after carbachol (n=13). Statistical significance was established by a paired t test (p<0.0001). **E.** Q value for non-resonator neurons before and after carbachol treatment. Q is estimated as the ratio of the peak resonant frequency to the frequency where the maximum impedance has decay to half its value. Statistical significance was established by a paired t test (p<0.0001). **F.** Sample impedance profile of a resonator cell before (continuous line) and after (dotted line) carbachol has been added to the bath. **G.** Peak Resonance Frequency of resonant cells before and after carbachol (n=10). Statistical significance was established by a paired t test (p=0.916). **H.** Q value for resonator neurons before and after carbachol treatment. Q is estimated as the ratio of the peak resonant frequency to the frequency where the maximum impedance has decay to half its value. Statistical significance was established by a paired t test (p=0.56).

After carbachol treatment, the impedance curves of non-resonators became indistinguishable from those of resonators (Figure 3A), suggesting that acetylcholine enhances the recruitment of voltage-gated channels in the non-resonant population, bringing their response in line with that of the resonant population. A two-way ANOVA revealed no significant main effect of Type of Cell (i.e., resonators before carbachol, carbachol-treated resonators, and carbachol-treated non-resonators) (F(1.881, 56.44) = 0.06182, P = 0.93), indicating that the three types of cells were not significantly different from each other. Additionally, the interaction between Frequency and Type of Cell was not significant (F(3.167, 27.08) = 1.170, P = 0.34), suggesting that this lack of difference between the types of cells was consistent across all frequencies analyzed. In further support, the additional neurons recorded within each slice after carbachol treatment were uniformly classified as resonators, with a higher preferred resonance frequency (Figure 3B), affirming that acetylcholine creates a homogeneous population of resonant neurons—a distinct shift from the two distinct populations observed under control conditions (Supplemental Figures 1B and 3C).

**Figure 3.**
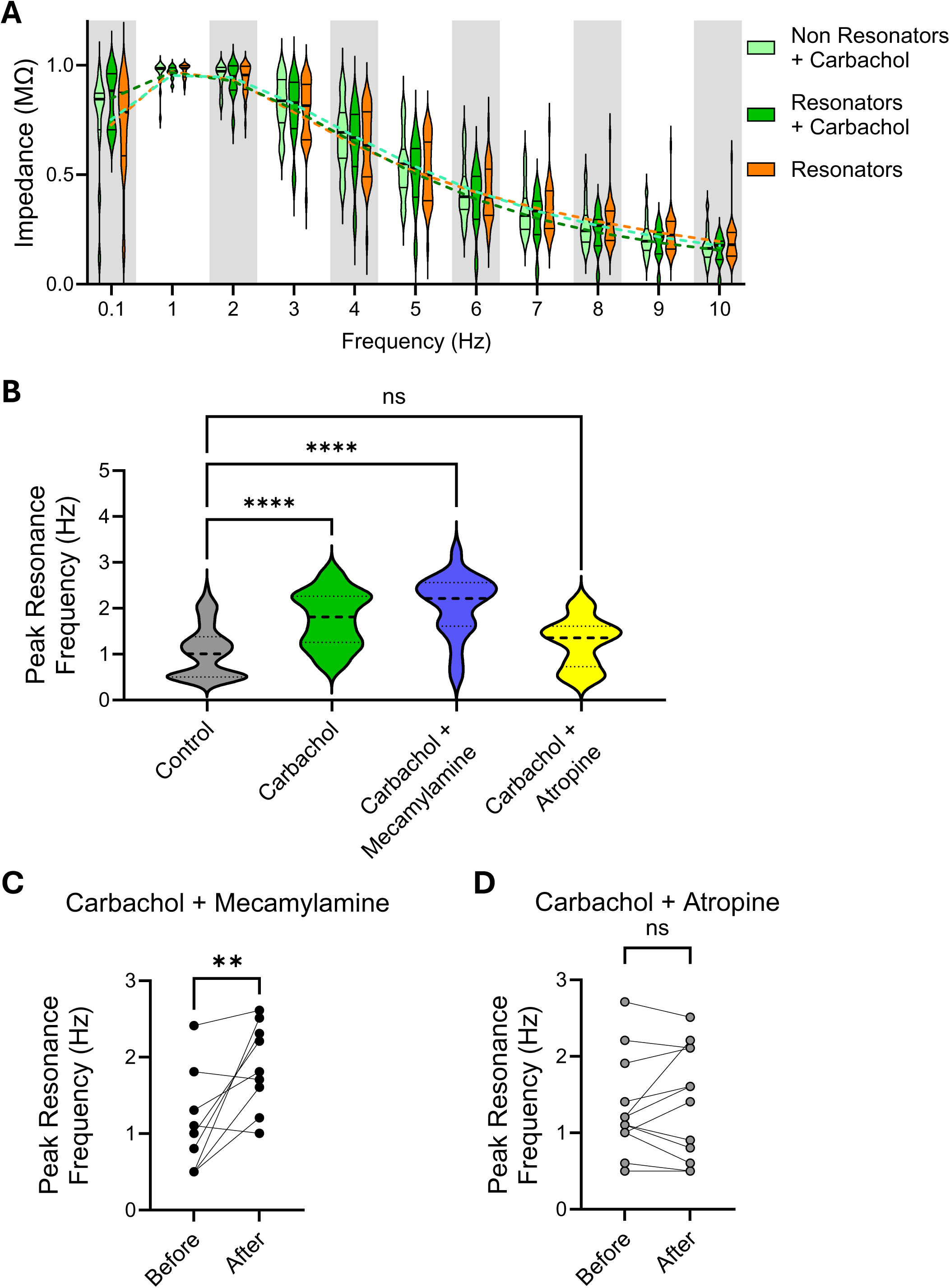
Metabotropic Acetylcholine Receptors Mediate Carbachol’s Effect. **A.** Rescaled values of impedance ([0 max]) from resonant cells at baseline, carbachol-treated resonant cells, carbachol-treated non-resonant cells binned around integer frequencies and compared using a 2-way ANOVA (Frequency x cell type: F (3.167, 27.08) = 1.170, p=0.34), followed by Tukey’s multiple comparisons test. **B.** Population data of the Preferred Resonance Frequency of all control, i.e. untreated neurons (n=72), and neurons from slices treated either with carbachol for at least 10 minutes (40 μM; n=33), carbachol plus mecamylamine, an acetylcholine ionotropic receptor blocker (10 μM; n=21), or carbachol plus atropine, an acetylcholine metabotropic blocker (1 μM; n=14). Statistical significance was determined using one-way ANOVA (F=22.32; p<0.0001) followed by Tukey multiple comparisons test. **C.** Preferred Resonance Frequency in the same neuron before and 10 minutes after treatment with carbachol plus mecamylamine (n=10). Statistical significance was determined using a paired t test (p=0.005). **D.** Preferred Resonance Frequency in the same neuron before and 10 minutes after treatment with carbachol plus atropine (n=13). Statistical significance was determined using a paired t test (p=0.47).

To discern which acetylcholine receptor is responsible for the observed effect on the preferred resonance frequency of monkey CA1 cells, we co-applied carbachol with either mecamylamine, a selective antagonist of ionotropic receptors, or atropine, an antagonist of metabotropic receptors. The increase in preferred resonance frequency caused by carbachol was blocked by atropine but was unmitigated by mecamylamine, indicating selective involvement of metabotropic receptors (Figure 3B). Similar effects of mecamylamine and atropine were observed in paired-recordings of the same neuron before and after the bath application of the drugs. This intra-cellular approach eliminates potential confounding variables that may arise when comparing different populations of neurons. In line with the results from the broader population of treated and untreated cells, co-application of carbachol with mecamylamine failed to prevent the change in the preferred resonance frequency (Figure 3C), while atropine prevented the change in the preferred resonance frequency when co-applied with carbachol. (Figure 3D).

These results indicate that activation of metabotropic acetylcholine receptors plays a crucial role in modulating the preferred resonance frequency of monkey CA1 neurons, highlighting the intricate regulatory mechanisms underlying cellular responses to cholinergic signaling.

## Cholinergic Regulation of Voltage-Gated Channels in Non-Resonators Neurons

We observed robust differences in the membrane’s voltage response to long hyperpolarizing current steps between the resonant and non-resonant populations, suggesting variations in voltage-dependent conductances, or the active properties of the membrane, across these two groups. Voltage-sensitive conductances shape the voltage response to time-varying inputs, enabling neurons to resonate. Hyperpolarization-activated Cyclic Nucleotide-gated (HCN) channels play a crucial role in the generation of rhythmic oscillations by producing an inward current, termed H-current, that manifests as a notable “sag” potential in response to long hyperpolarizing current steps ^18^.

We tested the active properties of monkey hippocampal neurons by applying a series of hyperpolarizing 1-second-long step currents and measuring the voltage response of the neurons (Figure 4A and D). These voltage measurements, along with all other experiments described here, were conducted in the absence of drugs that block synaptic inputs or action potentials. Although this approach resulted in noisier recordings, it preserved the natural cell physiology. The peak voltage of the ‘sag,’ measured early in the trace, was plotted against the injected current, and the slope of the voltage-current relationship was calculated (Figure 4B and E). To mitigate potential confounds due to variability across different cells, the slope of the ‘sag’ curve was normalized to that of a similar curve from voltage measurements at steady state. This intracellular normalization ensures consistent comparisons regardless of individual cell differences. We used the ratio of these slopes as an indicator of the amount of H-current present in the neurons.

**Figure 4.**
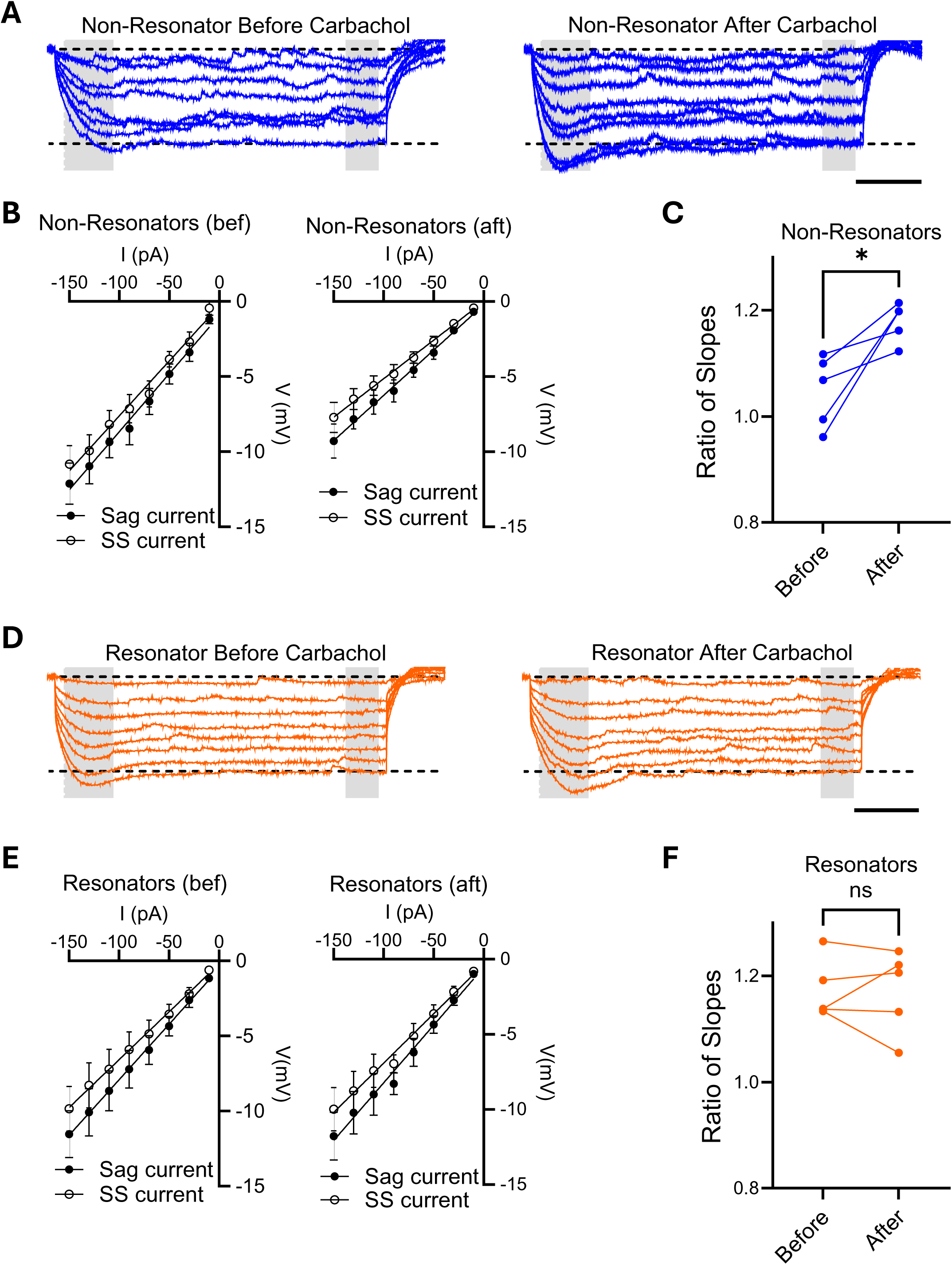
Modulation of Ih Current by Carbachol. **A.** Sample voltage traces of non-resonator neuron before (left) and after 10 minutes of carbachol been added to the bath (right). Step current pulses of 1 second and amplitudes varying from −150 pA to −10 pA were applied. The sag current was measured as the peak voltage between 50 ms and 150 ms after initiated the pulse (left gray area). The steady state current (SS) was measured as the average voltage of a 100 ms windows 900 ms after initiated the pulse (right gray area). Scale bar 200 ms. **B.** Relationship between current (I) and voltage (V) measured at the beginning of the pulse (sag) or at steady state (ss) in non-resonator cells, shown before (left) and after (right) the addition of carbachol to the bath. The slope of the sag current curves was normalized to the slope of the corresponding steady-state currents. **C.** Ratio of the slopes from I/V curves from non-resonant cells in B, before and after the addition of carbachol to the bath. **D.** Sample voltage traces of resonator neuron before (left) and after 10 minutes of carbachol been added to the bath (right). Step current pulses of 1 second and amplitudes varying from −150 pA to −10 pA were applied. The sag current was measured as the peak voltage between 50 ms and 150 ms after initiated the pulse (left gray area). The steady state current (SS) was measured as the average voltage of a 100 ms windows 900 ms after initiated the pulse (right gray area). Scale bar 200 ms. **E.** Relationship between current and voltage measured at the beginning of the pulse (sag) or at steady state (ss) in resonator cells, shown before (left) and after (right) the addition of carbachol to the bath. The slope of the sag current curves was normalized to the slope of the corresponding steady-state currents. **F.** Ratio of the slopes from I/V curves from resonant cells in E, before and after the addition of carbachol to the bath.

Neurons classified as non-resonators exhibited no “sag” voltage (Figure 4A, left, shaded area) and had a ratio of slopes around 1 (Figure 4B, left), signifying that an H-current is not detected in these neurons. We repeated the measurement in the same cell after adding carbachol to the bath and allowing it to incubate for at least 10 minutes. Carbachol rapidly increased the “sag” potential at hyperpolarizing steps (Figure 4A, right), with a concomitant increase in the ratio of slopes (Figure 4B and C), indicating an upregulation of H-currents.

Conversely, resonant neurons exhibited a “sag” response to hyperpolarizing potentials (Figure 4D, left, shaded area) and a ratio of slopes greater than 1 (Figure 4E, left), indicating the presence of H-currents in basal conditions. Adding carbachol to the bath and repeating the measurements in the same cell did not change the response to current steps (Figure 4D, right), resulting in similar I/V curves (Figure 4E, right) and similar ratios of slopes (Figure 4F).

To maximize the use of non-human primate slices, several other neurons were analyzed after the addition of carbachol to the bath. Figure 5 shows the resulting I/V curves for the population of neurons classified as non-resonators (Figure 5A), resonators (Figure 5B), and neurons treated with carbachol without established resonant classification (Figure 5C). The ratio of slopes for the population of resonant cells is significantly larger than that of non-resonant cells, indicating a correlation between the amount of H-current and the property of resonating at higher frequencies. Neurons from slices treated with carbachol for at least 10 minutes exhibited a ratio of slopes similar to that of resonant cells, further suggesting that carbachol homogenizes the population of hippocampal neurons in non-human primates (Figure 5D).

**Figure 5.**
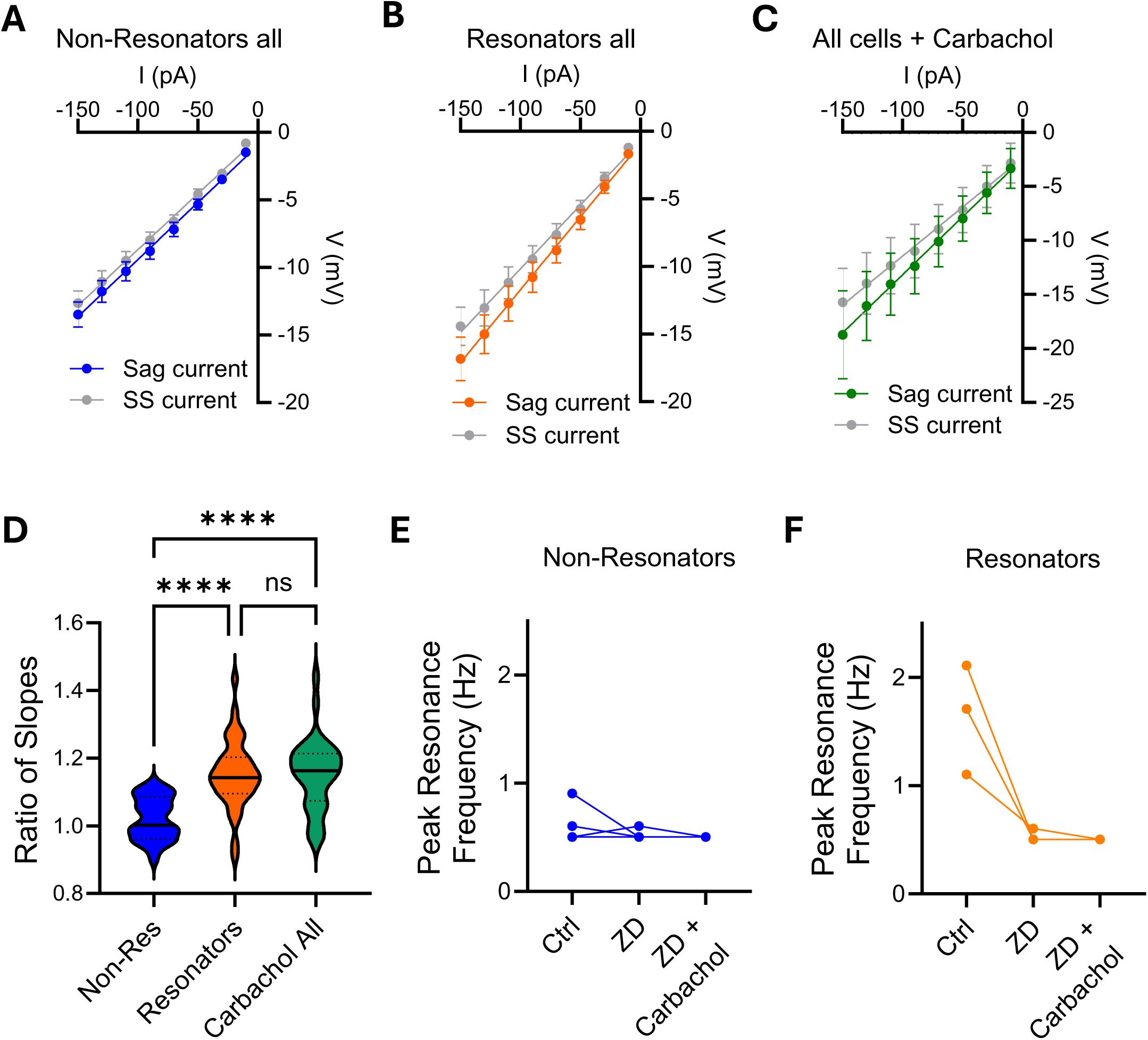
Population Data of Carbachol Effect on Ih Current. A-C. Relationship between current and voltage measured at the beginning of the pulse (sag) or at steady state (ss) in all cells classified as non-resonator cells (A; n=24), resonators (B; n=39), or all cells that has been treated with carbachol (C; n=33). **D.** The slope of the sag current curves was normalized to the slope of the corresponding steady-state currents. Statistical analysis was performed using one-way ANOVA (F=16.93, p<0.0001), followed by Tukey’s multiple comparison test. **E.** Preferred Resonance Frequency of non-resonator cells before and after treatment with ZD7288. Carbachol was then added to the bath in the continued presence of ZD7288. **F.** Preferred Resonance Frequency of resonator cells before and after treatment with ZD7288, demonstrating the dependence of resonance on HCN channels. Carbachol was then added to the bath in the continued presence of ZD7288.

All together these results suggest that the activation of metabotropic acetylcholine receptors can rapidly modulate the intrinsic properties of monkey hippocampal neurons via the upregulation of H-currents, thereby increasing the preferred resonant frequency.

Concurrent with this, blockade of HCN channels with ZD7288 eliminates the facilitatory effect that carbachol has on cellular resonance in non-resonant neurons (Figure 5E). Additionally, blockade of HCN channels disrupts the resonance of resonant neurons, indicating that HCN channels are a key mediator of cellular resonance across the population of CA1 pyramidal neurons in primates. More importantly, ZD7288 occludes any effect of carbachol on the preferred resonance frequency, indicating that the upregulation of HCN channel function is responsible for the observed modulation (Figure 5F).

## Discussion

Acetylcholine plays a key role in the function of the hippocampal memory system, evidenced by the encoding deficits caused by muscarinic antagonists ^17,24^ and the facilitatory effects of acetylcholinesterase inhibitors in early Alzheimer’s pathology ^25^. To date the link between the cellular effects of acetylcholine and the microcircuit computations supporting memory have remained elusive. Our results provide a mechanism by which acetylcholine acutely modulates cellular physiology within the hippocampal circuit – providing a means to rapidly shift the circuit into a theta-rhythmic state that could facilitate temporal coordination within the network. This is particularly critical in primates, where the transitory nature of theta oscillations suggests rapid neuromodulation of cellular properties that facilitate synchronization within hippocampal microcircuits.

Our findings reveal heterogeneity in the resonance properties of non-human primate hippocampal CA1 neurons. One population, the non-resonant neurons, displays an impedance profile similar to pink noise filtering, typical of the passive capacitive properties of the cytoplasmic membrane. The second population, the resonant neurons, exhibits lower impedance at low frequencies and a higher voltage response within theta band frequencies due to a heightened impedance profile that extends toward 4Hz. These resonant neurons demonstrate an increased contribution of voltage-gated channels, as indicated by a prominent sag potential, which is the hallmark of hyperpolarization-activated channels. In the presence of carbachol, these two distinct physiological phenotypes of CA1 pyramidal cells homogenize into a uniform theta resonant population with similar impedance profiles through a signaling pathway mediated by metabotropic acetylcholine receptors, which increases somatically recorded HCN channel activity. Through these experiments, we have gained mechanistic insight into the role acetylcholine has in hippocampal circuit dynamics and theta oscillations in primates.

By modulating the impedance profile of the neurons, acetylcholine may create a more uniform filtering profile, which in turn heightens depolarization in response to theta-range input and places the neuronal population in a state that is more permissive of theta oscillations. This swift modulation by acetylcholine suggests that afferents from the basal forebrain may facilitate the participation of these neurons in synchronized oscillations within hippocampal microcircuits. This mechanism may explain the episodic nature of theta oscillations in primates, contrasting with the persistent nature observed in rodents.

Oscillations that emerge at the network level are shaped by interacting mechanisms, each acting at different levels of organization: neuronal resonance ^22,26^, intrinsic circuit rhythmicity ^27^, and inherited rhythmicity from extrinsic input ^28^. The ability of each mechanism to drive network rhythmicity is determined by both the preponderance of each mechanism and its capacity to propagate across levels of organization ^29^. The rhythmicity manifested at each level of organization may be better framed as an interaction of low-pass and high-pass filtering ^30,31^, and therefore the rhythmic profile of oscillatory activity at the network level is the combined interaction of multiple layers of filtering ^29^. Engaging a single mechanism will not necessarily synchronize across levels to drive network oscillations, but it can change the propensity of the network toward engaging in oscillatory behavior ^32^.

Here we show that carbachol causes CA1 pyramidal cells to become a homogenous population with a uniform filtering profile that heightens depolarization to theta range input and attenuates the voltage response to sub-theta frequency input -- convergent with the cholinergic effect on theta in rodents ^33^. Though resonance profiles at the intracellular level might not directly dictate rhythmicity at the network level, homogeneity across the CA1 population increases the ability to conduct the theta resonant profile across levels, whereas heterogeneity creates a dissonance across the cellular filters that stymies the sensitivity to theta in the aggregate population. In essence, cholinergic input to CA1 puts the population in a state that is permissive to theta oscillations. By determining how acetylcholine shapes the responsive frequency range of hippocampal neurons, we have parameterized the most basal level of network-level rhythmicity.

Transiently entraining the hippocampal circuit with an oscillatory regiment could provide a unique mode of computation by creating a brief window of synchronized communication between entorhinal cortex and CA1. In rodents, each station within the canonical tri-synaptic circuit participates in a separable theta-rhythmic oscillator ^34^. Efficient communication across the hippocampal formation requires coherence of its layers – resulting in a neural code that is parcellated by the cadence of theta ^35,36^. Acetylcholine is hypothesized to inhibit intrinsic communication within the recurrent circuits of CA3 and CA1 while leaving the influence of entorhinal inputs on CA1 intact ^12,37,38^. Consequently, transient theta oscillations may correspond to a processing state that gives entorhinal input privileged access CA1, allowing it to acutely drive CA1 neurons and induce resultant plasticity. At a cognitivist level, this process may broadly correspond to pointedly attending to the experience of the external world, while disengaging from internal models derived from previous experience.

This study provides a foundation for bridging cellular physiology to hippocampal circuit function and provides mechanistic insight into the influence of acetylcholine on hippocampal circuit dynamics in primates, linking oscillations in the local field potential to neuropharmacology and behavioral states. *In vivo* studies will be important to determine whether acetylcholine is sufficient to drive network-level theta oscillations or if cholinergic-mediated cellular rhythmicity is only one component of primate hippocampal theta, providing mechanistic clarity in interpreting theta as a biomarker. Future studies should also delineate the specific cellular pathways mediating changes to membrane resonance in primates while also determining the factors shaping the transientness of the resonance and terminating theta wave bouts in the hippocampus of non-human primates. By developing a cellular-level understanding of hippocampal circuitry in primates, we gain the opportunity to learn what network mechanisms translate well across species and where primates are unique – providing guidance towards future research, refining targets for medical intervention, and uncovering the foundation of hippocampal processing in human cognition.

## Methods

All experiments were conducted in accordance with the animal care and use policies of the University of Washington and the Washington National Primate Research Center (WaNPRC). Hippocampi were collected from male and female Macaca nemestrina, aged 5 to 17 years, through the Tissue Distribution Program of the WaNPRC. The animals were transcardially perfused with a neuroprotective N-Methyl-D-glucamine artificial cerebrospinal fluid (NMDG-aCSF) solution during euthanization.

Immediately after dissection, hippocampi were placed in ice-cold, oxygenated NMDG-aCSF (93 mM NMDG; 93 mM HCl; 2.5 mM KCl; 1.2 mM NaH2PO4; 30 mM NaHCO3; 20 mM HEPES; 25 mM glucose; 5 mM Na-ascorbate; 2 mM thiourea; 3 mM Na-pyruvate; 10 mM MgSO4; 0.5 mM CaCl2). Hippocampal slices (300 µm thick) were then prepared using a Precisionary Instruments VF-200-0Z compresstome. Slices were recovered for 10 minutes at 32°C in oxygenated NMDG-aCSF, then transferred to room-temperature, oxygenated HEPES-based aCSF (92 mM NaCl; 2.5 mM KCl; 1.2 mM NaH2PO4; 30 mM NaHCO3; 20 mM HEPES; 25 mM glucose; 5 mM Na-ascorbate; 2 mM thiourea; 3 mM Na-pyruvate; 10 mM MgSO4; 0.5 mM CaCl2) for an additional 60 minutes of recovery.

Recordings were made at 25°C, 32°C, or 38°C, as indicated in the text, with slices submerged in recirculated, oxygenated aCSF (124 mM NaCl; 2.5 mM KCl; 1.2 mM NaH2PO4; 24 mM NaHCO3; 5 mM HEPES; 12.5 mM glucose; 2 mM MgSO4; 2 mM CaCl2). A K⁺-based internal solution (115 mM K-Gluconate; 20 mM KCl; 10 mM HEPES; 2.5 mM MgCl2; 4 mM Na2ATP; 0.4 mM Na3GTP; 10 mM Na-phosphocreatine; 0.6 mM EGTA) was used in electrodes with a resistance of 3-5 MΩ.

All recordings were made from the intermediate region of the longitudinal axis of the hippocampus. This portion of the body of the hippocampus extended from the coronal plane in-line with the posterior tip of the uncus to 4 mm further posterior. Neurons for recording were visually identified within the CA1 region, which was conservatively identified as the portion of the ammonic field that was just lateral to a horizontal line extending from the hippocampal fissure. Series and membrane resistance were monitored to ensure the quality of whole-cell recordings. Cells with a resting membrane potential less negative than −55 mV were discarded. Recordings were performed using MultiClamp and Clampfit software (Axon Instruments). Unless otherwise noted in the text, no additional toxins were added to the bath. The drugs used were Carbachol (USP, 40 µM), atropine (Millipore-Sigma, 1 µM), mecamylamine (Tocris, 10 µM), XE991 (Tocris, 10 µM), and ZD7288 (Tocris, 10 µM).

Current injections included step protocols and custom ‘ZAP’ currents, which consisted of a sinusoid of constant amplitude (25, 50, or 100 pA) that varied in frequency from 1 to 20 Hz. The largest amplitude ZAP that did not induce action potentials was used, and traces where action potentials were elicited were discarded. In a subset of cells, neurobiotin was included in the patching pipette, and the slices were fixed and stained post-hoc using DAB (Vector Labs). Data were analyzed using Clampex (Axon Instruments) and Igor Pro (WaveMetrics).

## Acknowledgements

This work was funded by R21 MH 126126-01A1 (AB and JR) and WaNPRC/ITHS Ignition Award (AG and JR).

## Authors contributions

A.G., J.R., and A.B. conceived the project and designed the experiments. A.G. and J.R. conducted the experiments and collected data. A.G., J.R., and A.B. performed data analysis. A.B. and J.R. wrote the manuscript.

---

**Supplemental Figure 1.**
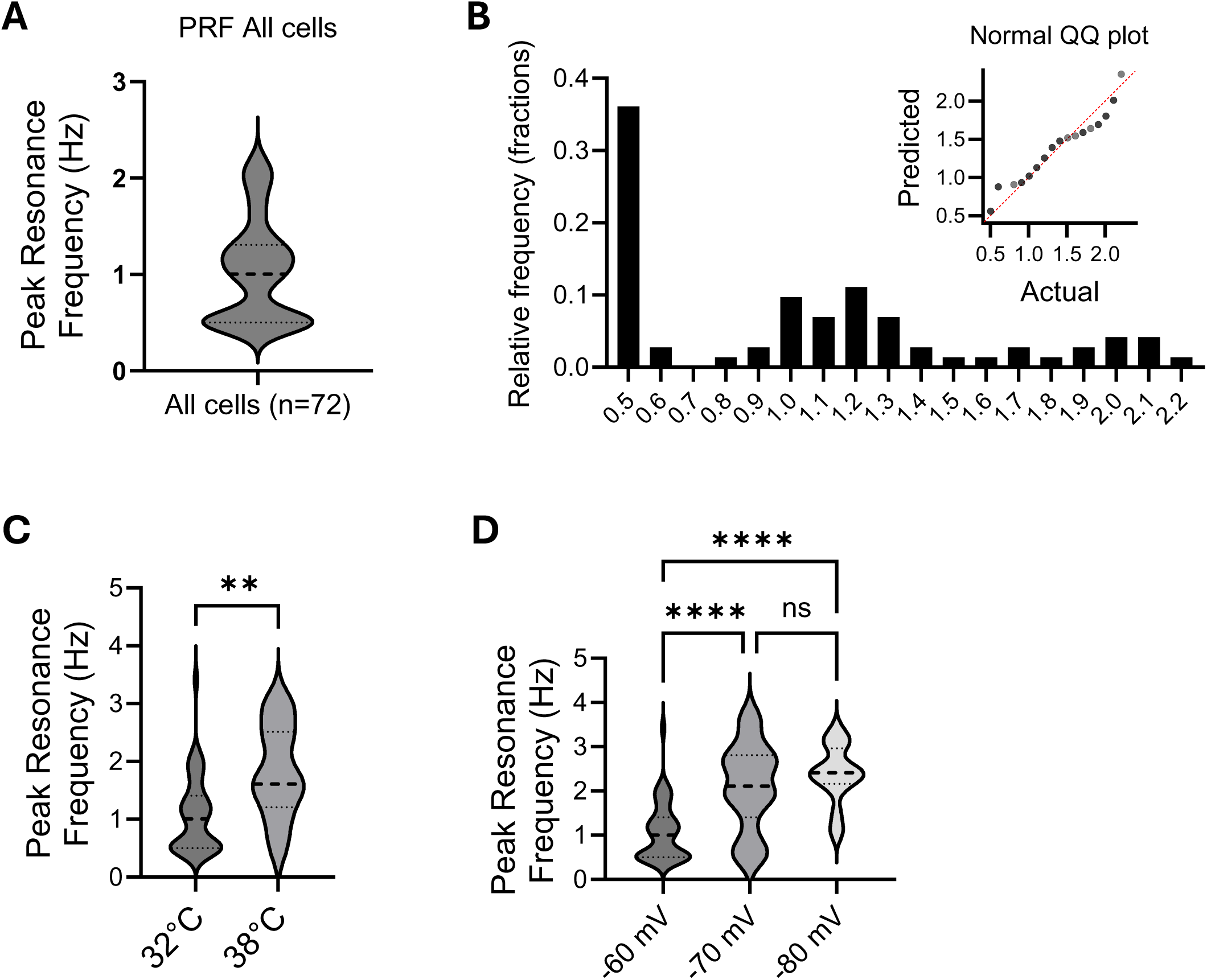
**A.** Violin plot showing the Preferred Resonance Frequency of all neurons recorded. **B.** Histogram of the Preferred Resonance Frequency of all neurons recorded as in A. Inset, QQ plot comparing the distribution of the dataset in A to a theoretical normal distribution. **C.** Preferred Resonance Frequency of CA1 neurons recorded at different temperatures as indicated. Two tailed unpaired t test, p=0.0025 **D.** Preferred Resonance Frequency of CA1 neurons in current clamp at different baseline voltages, as indicated. A one-way ANOVA revealed a significant difference (F = 23.81, p < 0.0001), followed by Tukey’s multiple comparison test.

**Supplemental Figure 2.**
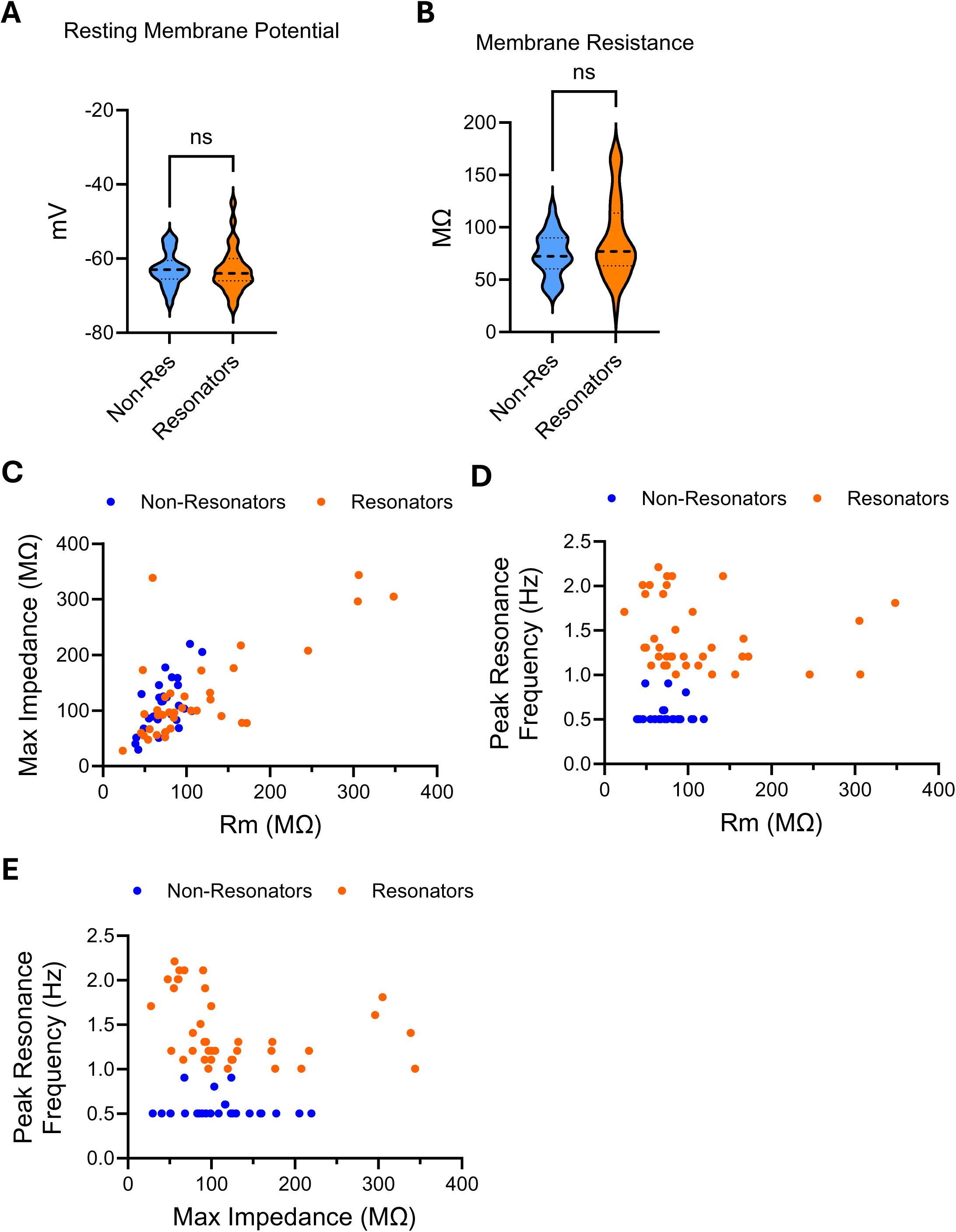
**A.** Violin plot displaying the Resting Membrane Potential of non-resonator and resonator cells. A two-tailed Mann-Whitney test revealed no significant difference between the groups (p = 0.5958). **B.** Violin plot displaying the membrane resistance of non-resonator and resonator cells. A two-tailed Mann-Whitney test revealed no significant difference between the groups (p = 0.2816). **C.** Scatter plot showing the relationship between membrane resistance and maximal impedance in non-resonator and resonator cells. **D.** Scatter plot showing the relationship between membrane resistance and Preferred Resonance Frequency in non-resonator and resonator cells. **E.** Scatter plot showing the relationship between maximal impedance and Preferred Resonance Frequency in non-resonator and resonator cells.

**Supplemental Figure 3.**
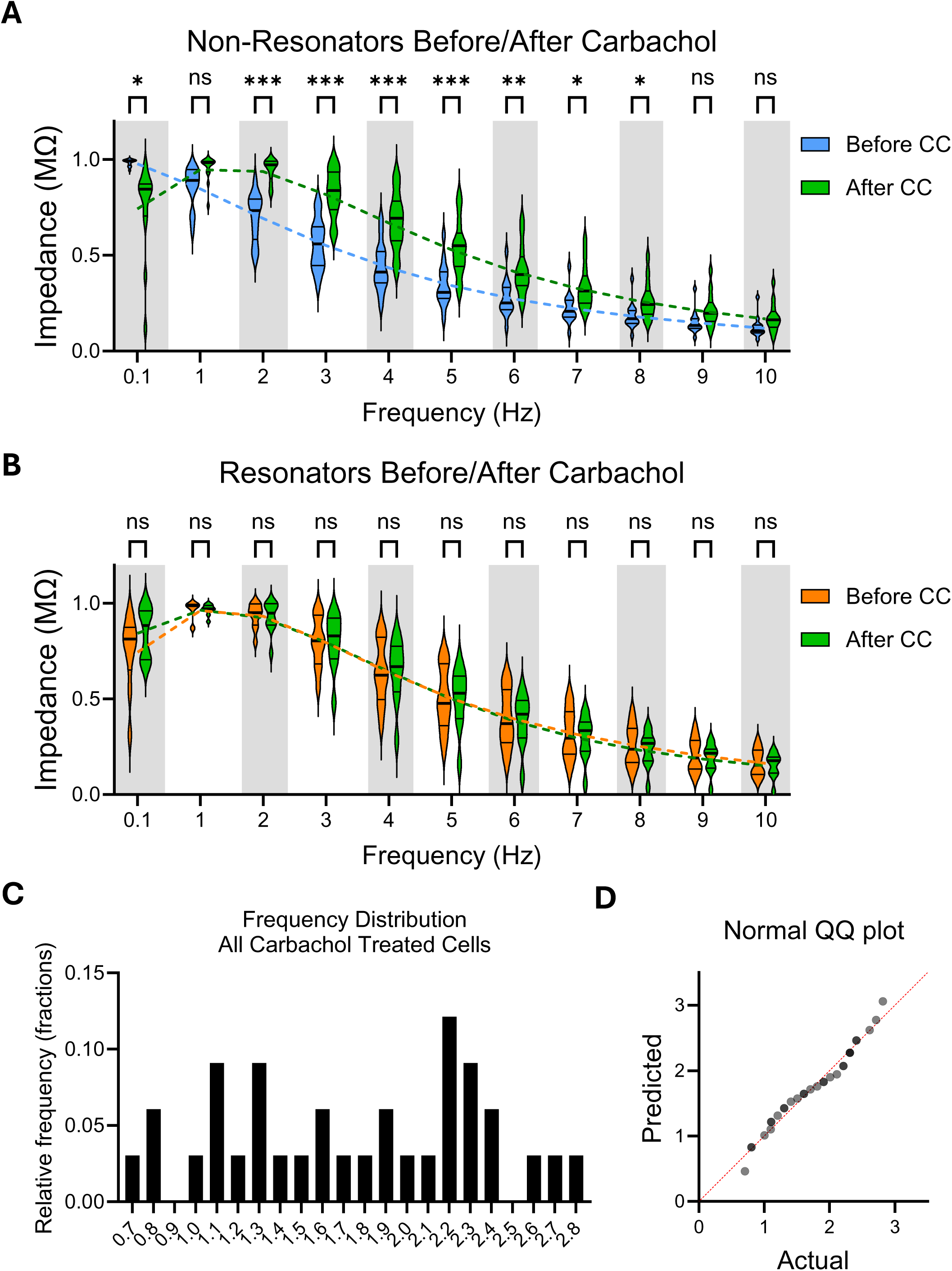
**A.** Rescaled values of impedance ([0 max]) from non-resonator cells before and after treatment with carbachol and binned around integer frequencies and compared using a 2-way ANOVA (Frequency x before/after carbachol: F (1.718, 20.62) = 23.85, p<0.001), followed by Tukey’s multiple comparisons test. **B.** Rescaled values of impedance ([0 max]) from resonator cells before and after treatment with carbachol and binned around integer frequencies and compared using a 2-way ANOVA (Frequency x before/after carbachol: F (1.328, 11.95) = 1.353, p=0.28), followed by Tukey’s multiple comparisons test. **C.** Histogram of the Preferred Resonance Frequency of all neurons treated with carbachol. **D.** QQ plot comparing the distribution of the dataset in C to a theoretical normal distribution

